# Genomic deletions and rearrangements in monkeypox virus from the 2022 outbreak, USA

**DOI:** 10.1101/2022.09.16.508251

**Authors:** Crystal M. Gigante, Matthew Plumb, Ali Ruprecht, Hui Zhao, Vaughn Wicker, Kimberly Wilkins, Audrey Matheny, Theodora Khan, Whitni Davidson, Mili Sheth, Alex Burgin, Mark Burroughs, Jasmine Padilla, Justin S. Lee, Dhwani Batra, Ethan E. Hetrick, Dakota T. Howard, Jacob Garfin, Lisa Tate, Shan J. Hubsmith, Rafael M. Mendoza, Danielle Stanek, Sarah Gillani, Michelle Lee, Anil Mangla, David Blythe, Sean SierraPatev, Kristin Carpenter-Azevedo, Richard C. Huard, Glen Gallagher, Joshua Hall, Stephanie Ash, Lynsey Kovar, Matthew H. Seabolt, Michael R. Weigand, Inger Damon, Panayampalli S. Satheshkumar, Andrea M. McCollum, Christina L. Hutson, Xiong Wang, Yu Li

**Author notes:** The findings and conclusions in this report are those of the author(s) and do not necessarily represent the views of the Centers for Disease Control and Prevention.

## Abstract

Genomic surveillance of monkeypox virus (MPXV) during the 2022 outbreak has been mainly focused on single nucleotide polymorphism (SNP) changes. DNA viruses, including MPXV, have a lower SNP mutation rate than RNA viruses due to higher fidelity replication machinery. We identified a large genomic rearrangement in a MPXV sequence from a 2022 case in the state of Minnesota (MN), USA, from an abnormal, uneven MPXV read mapping coverage profile in whole-genome sequencing (WGS) data. We further screened WGS data of 206 U.S. MPXV samples and found seven (3.4 percent) sequenced genomes contained similar abnormal read coverage profiles that suggested putative large deletions or genomic rearrangements. Here, we present three MPXV genomes containing deletions ranging from 2.3 to 15 kb and four genomes containing more complex rearrangements. Five genomic changes were each only seen in one sample, but two sequences from linked cases shared an identical 2.3 kb deletion in the 3’ terminal region. All samples were positive using VAC1 and Clade II (formerly West African)-specific MPXV diagnostic tests; however, large deletions and genomic rearrangements like the ones reported here have the potential to result in viruses in which the target of a PCR diagnostic test is deleted. The emergence of genomic rearrangements during the outbreak may have public health implications and highlight the importance of continued genomic surveillance.

---

A global effort has been made to sequence monkeypox virus (MPXV) associated with the current outbreak to track virus lineages and mutations. All sequences associated with the outbreak in non-endemic countries have belonged to Clade IIb (formerly named West African MPXV found east of the Dahomey Gap) [1, 2]. Two lineages within Clade IIb have been identified in 2022, one predominant lineage (B.1) and a second more diverse lineage associated with a handful of cases with links to West Africa or the Middle East (A.2). Among lineage B.1 MPXV, there have been few single nucleotide polymorphisms (SNPs) observed thus far compared to RNA viruses such as influenza virus and SARS-CoV-2 [3, 4], mostly due to the lower mutation rate of higher fidelity replication machinery in MPXV. In fact, most mutations that have been observed in the current outbreak genomes have been attributed to host APOBEC3 activity [3-5].

It is believed that poxviruses compensate for a relatively low rate of point mutations by exhibiting frequent genomic duplications, losses, and gains by recombination and horizontal gene transfer [6-10]. Genomic rearrangements generate diversity in viral populations and can play an important role in the evolution of viruses. Indeed, genomic duplications and deletions that affect viral pathogenicity have been reported for vaccinia virus in culture and under selection conditions [11, 12]. Only one previous large genomic rearrangement has been reported in a Clade I MPXV from Sudan in 2005 (KC257459) [13]. Quick and accurate identification of large genomic changes can be challenging, particularly for short-read sequencing platforms widely used in public health laboratories, yet such mutations can cause very rapid phenotypic changes to the virus.

We observed an unusual read mapping profile in MPXV whole-genome sequencing data from a lesion specimen derived from a case-patient, MN0001. Mapping reads to 2022 MPXV lineage B.1 reference sequence MA001 (ON563414.3) produced an absence of mapped reads for a large region towards the right (3□) end of the genome and an overabundance of reads on the left-hand side of the genome (5□ end). The result prompted us to revisit sequencing data of 206 U.S. MPXV positive samples and identify a total of seven (3.4%) genomes (including MN0001) with similar abnormal read coverage profiles (Figure 1). All seven genomes contained gaps in the read mapping profiles, and four genomes also contained high coverage plateaus, all of which were observed near the ends and not in the middle of the genomes.

**Figure 1.**
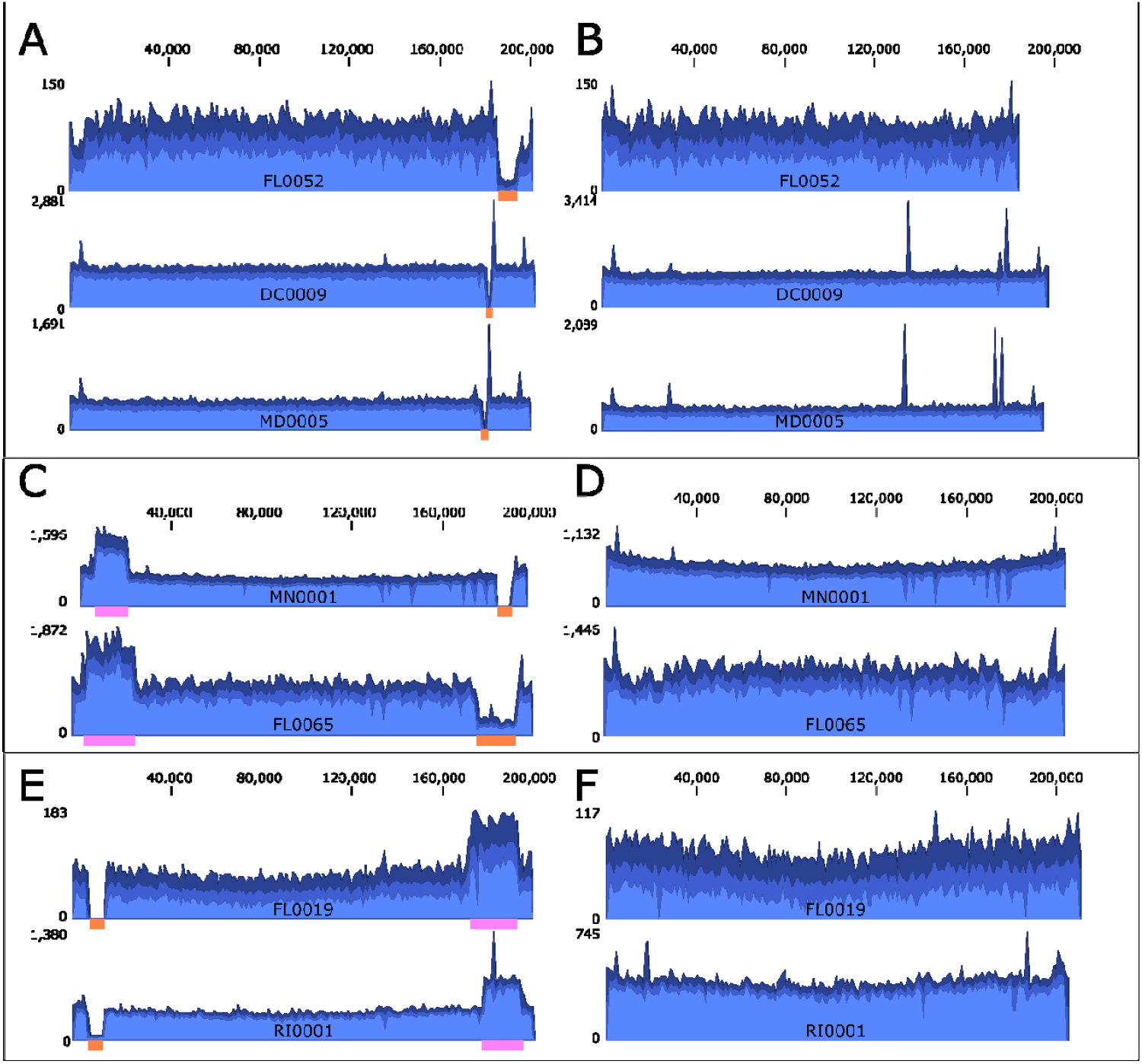
Indication of genomic rearrangement by examination of read mapping. A. – F. Coverage graph of Illumina short read data mapped to MPXV lineage B.1 MA001 reference genome (A, C, E) or final genomes containing proposed genomic rearrangement or deletion (B, D, F). Orange bars indicate regions of genomes with no or low coverage, and pink bars mark regions with high coverage. MPXVs with genomic deletions are shown in top box; MPXVs with right terminus rearrangement are shown in center box; and MPXVs with left terminus rearrangement are shown in bottom box. Read depth is shown in blue with scale bar to the left. Nucleotide alignment position is shown above.

Close investigation of reads mapped near the boundaries of observed coverage gaps revealed two types of putative genomic rearrangement. In three sequences (DC0009, MD0005, and FL0052), reads spanned the coverage gap, suggesting a deletion in the genome. The putative deletion was confirmed by mapping reads to a reference-based assembly genome in which the sequence that spanned the gap in the reference was removed. Multiple reads spanned the junction and reads mapped more evenly across the draft genome that contained the deletion (Figure 1B).

These data revealed three genomic deletions ranging from 2.3 to 15 kb compared to reference genome MA001 (Table 1). Interestingly, viral sequences from two cases (MD0005 from Maryland and DC0009 from Washington D.C.) contained identical deletions of 2.3 kb in the right terminus of the genome that resulted in putative truncation of B17R and deletion of B18R coding sequences (CDS) (Figure 2). The deletion observed in FL0052 was much larger (>14 kb) and involved deletion of eight CDS and putative truncation of B22R. This deletion also shortened the inverted terminal repeat (ITR) region to 784 bases for this strain (Figure 2), which is similar to the ITR length seen in Variola virus.

**Table 1.** Genome details for seven 2022 MXPV from U.S. with genomic rearrangements. Del: deletion; ins: insertion. List of genes deleted and duplicated in each case can be found in Table S1.

| Sample ID | Genome size | Rearrangement | Terminus | Terminal region changes |
| --- | --- | --- | --- | --- |
| MN0001 | 204181 bp | Insertion-deletion | Right | 13.4 kb del, 20.7 kb inverted ins from left |
| RI0001 | 205900 bp | Insertion-deletion | Left | 13 kb del, 21.7 kb inverted ins from right |
| DC0009 | 194855 bp | Deletion | Right | 2.3 kb del |
| MD0005 | 194851 bp | Deletion | Right | 2.3 kb del |
| FL0019 | 210731 bp | Insertion-deletion | Left | 14.1 kb del, 27.7 kb inverted ins from right |
| FL0052 | 182811 bp | Deletion | Right | 14 kb del |
| FL0065 | 199828 bp | Insertion-deletion | Right | 23.8 kb del, 26.4 kb inverted ins from left |

**Figure 2.**
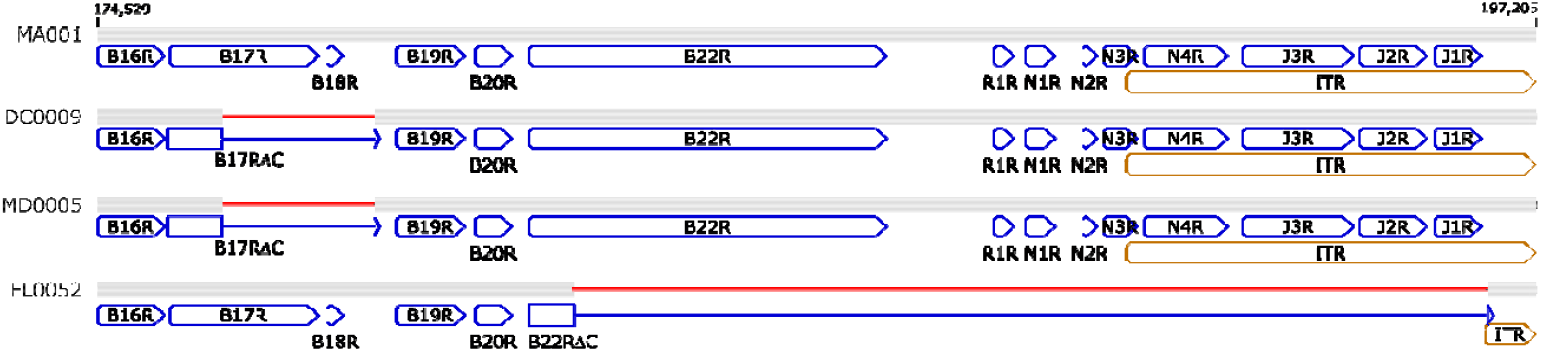
Position of genomic deletions in DC0009, MD0005 and FL0052. Graphical depiction of the alignment of right genomic termini of DC0009, MD0005 and FL0052 to MA001 reference genome. FL0052 contained a deletion of J2R (CrmB, soluble TNF receptor), which is the target of MPXV generic and MPXV Clade II specific PCR assays; however, this CDS and three other deleted ITR CDS were intact in the 5□ end of the genome (not shown). Deletions relative to MA001 are highlighted in red in the right terminus. Annotations are depicted in blue. Gray bars indicate identity, and black bars indicate differences with MA001. Nucleotide positions are relative to MA001.

The remaining four sequences (MN0001, RI0001, FL0019 and FL0065) contained a gap in read coverage as well as a high coverage plateau on the end of the genome opposite the gap. For these sequences, no reads spanned the gap, suggesting a more complex rearrangement rather than a simple deletion. When reads were mapped back to draft genomes containing deletions corresponding to the coverage gap, all reads in the putative junction area mapped only to one side of the junction. Frequently, only half of the read sequence aligned to the MPXV genome at the junction while the other half differed from the reference (Figure S1). Manual investigation of these reads bordering the gap (Text S1) revealed sequence corresponding to both the right and left side of monkeypox virus genome sequences based on BLASTn alignment of individual reads. For example, MN0001 reads on the left side of the gap corresponded to genomic positions ~20,600 and ~183,500 in MA001, while reads from the right side of the gap corresponded to genomic positions ~6500 and ~191,000. These data suggest a genomic rearrangement in which one genomic terminus was deleted and replaced by an inverted duplication of the other terminus. We identified the prospective rearrangement junction locations by searching for the sequences from the reads bordering the gap in the draft genome. Once a potential rearrangement junction was identified, a new draft genome was produced by deleting from the gap to the proximal end of the genome. Then the new terminus was copied from the distal end of the reference-based consensus genome starting at the junction and pasted at the gap as reverse complement. Reads were then mapped to this new draft genome to confirm the rearrangement. Read mapping revealed even coverage across these genomes and manual inspection revealed reads evenly spanning the junction (Figure 1E – F).

In FL0019 and RI0001, the left terminus was deleted and replaced by sequence from the right terminus (Figure 3). FL0019 contained deletion of seven CDS, truncation of D9L, and duplication of 16 CDS. RI0001 exhibited deletion of six CDS and duplication of eight CDS. FL0065 exhibited deletion of seven CDS and duplication of 21 CDS.

**Figure 3.**
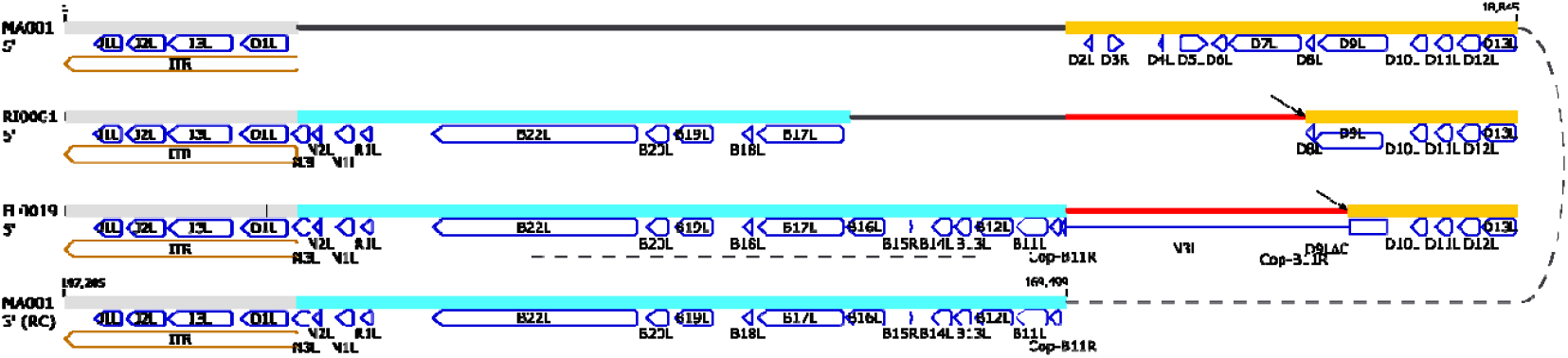
Large insertion-deletion in the left (5□) genomic terminus of RI0001 and FL0019. Graphical depiction of alignment of RI0001 and FL0019 to the 5□ and 3□ (RC: reverse complement) termini of MPXV lineage B.1 MA001 reference genome. Rearrangement junctions are highlighted by arrows; all sequences after junctions are identical to inverted 3□ genome terminus. Deletions relative to MA001 are highlighted in red; sequence from the 5□ terminus are shown in cyan; sequence from the 3□ terminus are shown in yellow; ITR sequence is shown in gray, differences in sequence relative to MA001 are shown in black. Annotations are depicted in blue below the sequence; gene name ending “L” or “R” refer to the direction of the ORF. Positions are relative to MA001. The 5□ and 3□ ends of MA001 are connected by a dotted line.

In MN0001 and FL0065, the right terminus was deleted and replaced by sequence from the left terminus (Figure 4). MN0001 exhibited deletion of four CDS and duplication of 15 CDS. The breakpoint occurred within the B22R CDS and P2L CDS for MN0001, resulting in a protein containing the N-terminal 801 aa from B21/B22L followed by the C-terminal 60 aa from P2L (Figure S2).

**Figure 4.**
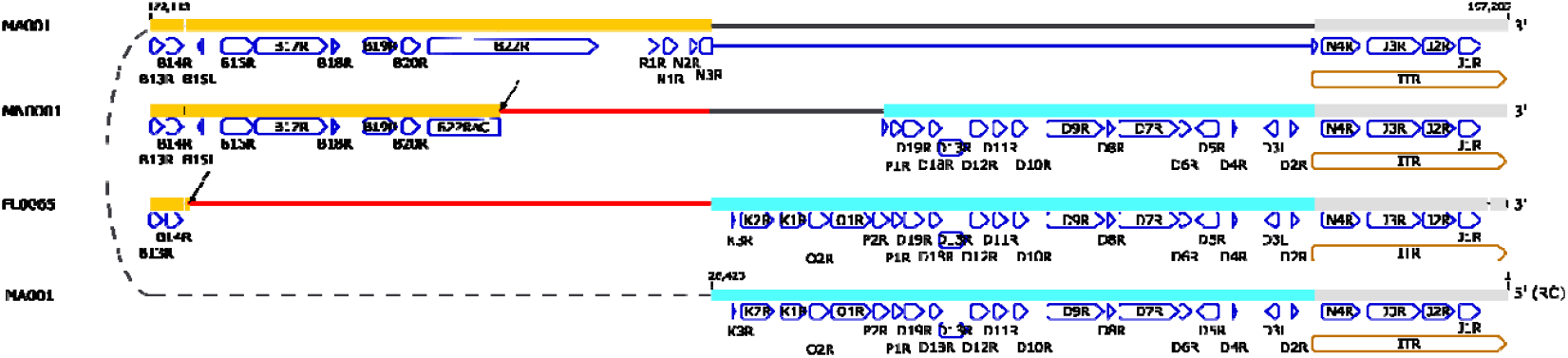
Large insertion-deletion in the right (3□) genomic terminus of MN0001 and FL0065. Graphical depiction of alignment of MN0001 and FL0065 to the 5□ (RC: reverse complement) and 3□ termini of MPXV lineage B.1 MA001 reference genome. Rearrangement junctions are highlighted by lightning bolts; all sequences after junctions are identical to inverted 5□ genomic terminus. Deletions relative to MA001 are highlighted in red; sequence from the 5□ terminus are shown in cyan; sequence from the 3□ terminus are shown in yellow; ITR sequence is shown in gray, differences in sequence relative to MA001 are shown in black. Annotations are depicted in blue below the sequence; gene name ending “L” or “R” refer to the direction of the ORF. Positions are relative to MA001. The 5□ and 3□ ends of MA001 are connected by a dotted line.

All seven samples were sequenced using metagenomics methods on both Illumina (shown in Figure 1) and Oxford Nanopore platforms. MN0001 and RI0001 were sequenced by Illumina sequencing using both metagenomics and tiled amplicon PCR-based approaches at two institutions. Similar read mapping profiles were observed with Oxford Nanopore data, but the tiled amplicon approach failed to yield obvious gaps and high coverage regions in RI0001. The presence of the junctions for all seven sequences were confirmed independently by short and long read sequence data. Lastly, PCR using primers designed to distinguish between the presence and absence of the rearrangement resulted in amplification across the rearranged junction for all seven sequences but not for other samples from the 2022 outbreak. Sanger sequencing of these amplicons confirmed the sequence of the junction (Text S2).

The presence of these terminal swapping rearrangements is supported by the read mapping profiles (Figure 1). The gaps in coverage reveal regions of the genome that were deleted; and the high coverage regions at the other terminus correspond to the duplicated genomic terminus. One factor that may confuse interpretation of the read mapping profiles is the appearance of the ITR region on the deleted terminus. Because this sequence is identical on both termini in the reference, many read mappers will randomly assign reads to both termini, producing relatively even coverage.

Identification and assembly of genomes containing terminal swapping rearrangements was not straight forward. Attempts at reference-based consensus sequence generation using either Illumina or nanopore reads produced incorrect genomes containing simple deletions and no insertion. *De novo* assembly with only short reads was unable to span the junction region due to failure to resolve identical genomic termini. This is a problem for assembly of all poxvirus genomes in the ITR region. Visualization of assembly graphs from SPAdes or Unicycler revealed the insertion region containing the ITR could be assembled in the reverse complement to either end of the main contig (Figure S3). *De novo* assembly with long reads using SPAdes or Unicycler was able to produce more complete contigs, including contigs that crossed the junction confirming the genomic rearrangement. Likewise, hybrid assembly methods using SPAdes or Unicycler were successful at producing contigs that crossed the junctions; however, in all cases, some manual assembly of the genomes was required. The existence of MPXV genome rearrangement activities indicate that an optimized WGS wet lab strategy and bioinformatics solutions are needed for high-throughput testing for MPXV genomic surveillance purposes.

RI0001, FL0052, and FL0065 added additional complications to genome assembly, as these samples appeared to contain mixed population of virus, with some sequence reads mapping evenly across reference genome MA001. The ratio of rearranged to wildtype virus varied for each sample from 10:1 for FL0052, 7:1 for RI0001, and 3:1 FL0065, based on read depth. In each instance, the majority of reads supported the genomic rearrangement. Each sample was sequenced three times (Illumina metagenomic, ONT metagenomic, and PCR Sanger), and the presence of mixed populations was confirmed by each method. The presence of two mixed genomes could indicate the MPXV genome was changing in these patients, but only partially. However, we cannot rule out the possibility that the sample was contaminated. The presence of mixed populations further complicated genome assembly, and these genomes were assembled manually. *De novo* assembly produced contigs that corresponded to both genomes: with and without rearrangement.

All seven cases associated with these MXPV sequences were male ranging from 30 to 42 years of age with symptom onset from June 8 to June 27 (Table 2). Five cases had lesions localized to genitals, perianal region, and/or mouth, and two cases had widespread lesions that spread to the trunk. MN0001, DC0009, and MD0005 had traveled to Europe during the estimated incubation period. The remaining four cases only reported travel within the USA. RI0001 reported sexual contact with four persons, two of which were long-term partners from MA. Sequences from the MA cases did not have evidence of the rearrangement observed in RI0001; however, they were identical to the minor (not rearranged) virus in RI0001. Follow-up with the other two reported contacts was unsuccessful. The FL0019 case reported a likely exposure at an event associated with seven additional monkeypox cases, all likely linked to a single index case. None of the MPXV sequences from those eight cases showed evidence of the rearrangement seen in FL0019. MPXV sequences from DC0009 and MD0005 contained an identical 2.3 kb deletion. The cases associated with these MPXV sequences were long-term partners with a common sexual encounter with a confirmed monkeypox positive case in Europe (Table 2). Interestingly, two SNPs were observed between the whole genome sequences of MD0005 and DC0009.

**Table 2.** Case details for seven associated 2022 MXPV from U.S. with genomic rearrangements. Travel indicates locations visited during incubation period or location of likely exposure. Estimated number and location of lesions are indicated. Localized: lesions localized to genitals, perianal area, or face/lip. Widespread: lesions spread to trunk. Known monkeypox positive contacts with sequencing data are indicated in contacts column. None: no reported high-risk contacts after symptom onset. Rearrangement in contacts was based on known, confirmed contacts from which MPXV sequences were available. TPOXX administration and hospitalization are indicated for each case. The first two letters in the Sample ID indicate the U.S. jurisdiction where the monkeypox case was confirmed: MN: Minnesota, RI: Rhode Island; MA: Massachusetts; DC: Washington D.C.; MD: Maryland; FL: Florida.

| Sample ID | Travel | Lesions | Known MPX Contacts | Rearrangement in contacts | TPOXX | Hospitalization |
| --- | --- | --- | --- | --- | --- | --- |
| MN0001 | Europe | <10, localized | None | Unknown | No | No |
| RI0001 | RI | widespread | Two MA cases | No | Yes | Yes |
| DC0009 | Europe | localized | MD0005 | Yes | No | No |
| MD0005 | Europe | localized | DC0009 | Yes | No | No |
| FL0019 | FL | >20, localized | FL case | No | Yes | No |
| FL0052 | FL | <20, widespread | None | Unknown | Yes | No |
| FL0065 | FL | <10, localized | None | Unknown | Yes | No |

This is one of the first reports describing large-scale genomic changes in MPXV during the 2022 outbreak. Given our observations for seven genomes out of 200, we suspect there may be other MPXV strains with similar rearrangements that may not have been properly identified. Thus, we provide a description of how to investigate and confirm the presence of putative large genomic deletions or terminal insertion-deletions based on sequence mapping. Automation of these observations based on calculated read coverage could help flag samples for manual review, long read sequencing or manual assembly.

Only two previous genomic rearrangements have been reported previously in MPXV. An 856 bp translocation has been described in a MPXV from Germany in 2022, that resulted in duplication of one gene and deletion of two CDS, and truncation of two CDS (https://www.biorxiv.org/content/10.1101/2022.07.23.501239v1.full.pdf). A larger genomic translocation was reported in a Clade I MPXV from Sudan in 2005 (KC257459) [13]. In that sequence, the deletion included most of the NMDA receptor CDS through most of 4^th^ CDS. The inverted insertion in the Sudan MXPV included 10.8 kb containing almost 12 CDS (MPXV_Zaire_1979-005 ORFs 5 – 16). The genomic location of this rearrangement is very similar to what we observed in MN0001 and FL0065. Indeed, among the eight 2022 MPXV described here and in preprint (https://www.biorxiv.org/content/10.1101/2022.07.23.501239v1.full.pdf) and the 2005 Sudan MPXV, several of the CDS involved are the same (Figures 3 and 4, Table S1), indicating the possibility that there may be a common mechanism. However, most of the exact breakpoints are unique, suggesting further investigation into the mechanism behind this genome terminal insertions/deletions is needed. Deletions in poxvirus genomes have been reported more frequently, including in a Clade II MPXV genome from Bayelsa (GenBank MT903341) [14]. Given the rarity of reported MPXV genomes with these types of genomic rearrangements, we performed sequencing using several different methods and confirmed results by PCR and Sanger sequencing; however, we cannot completely rule out the possibly of unanticipated artifacts caused by library preparation, improper sequencing read mapping or assembly.

Across orthopoxviruses, fixed recombinant events, gene loss, and gene duplication occur at a much higher rate at genomic termini, where non-essential and virulence genes are located. The ITRs are hot spots for large deletions and duplications, possibly through imperfect matching of repeat regions [15-20]. All deletions and genomic rearrangements observed in the 2022 MPXV genomes were located at the ends of the genomes. Deletions, gene duplications, and genomic rearrangements in the terminal regions have been associated with differences in poxvirus pathogenicity and host range [11, 16, 18, 21]. The seven genomic rearrangements we describe involved deletion and duplication of several genes that have been associated with poxvirus pathogenicity and host range (Table S1). It is difficult to speculate how these genomic changes might have affected viral pathogenicity, especially given the complex combination of deletion and insertion of several pathogenesis genes seen in some genomes. *In vitro* and animal studies are needed to better understand how these types of change affect viral pathogenesis and if duplication of genes may compensate for loss of others. It has been hypothesized that gene duplication and deletion in poxviruses may help them evolve to new host or selection conditions [7, 11]. Adaptive mutations that become fixed in a population over time may indicate MPXV is adapting to a new host (humans). It is currently unclear if these mutations are adaptive, neutral, or disadvantageous for the virus. We currently do not have evidence that MPXV with these specific genomic rearrangements are spreading and have only detected one genome with each rearrangement (with the exception of MD/DC cases).

None of the genomic rearrangements we describe involve the target regions for Orthopoxvirus PCR diagnostic tests, including the CDC FDA cleared non-variola orthopoxvirus test (VAC1) or an orthopoxvirus laboratory-developed test used by CDC (OPX3) [22]. Both VAC1 and OPX3 target essential genes in the center of the MPXV genome, where rearrangements and deletions are much less likely. None of the observed genomes contained disruptions or duplications of the F13L gene (encodes the target of TPOXX, an antiviral drug used for treatment of orthopoxvirus infections) and are not expected to affect the efficacy of TPOXX. One of the two copies of the target of Clade II MPXV-specific and MPXV generic real-time PCR assay (J2R/L, CrmB gene, Tumor necrosis factor (TNF) gene homolog) was deleted in FL0052. The copy on the 5□ end of the genome remained intact, and the sample was positive by Clade II MPXV-specific real-time PCR assay [23]. While we did not observe changes in the performance of diagnostic tests for these seven cases, large deletions and genomic rearrangements like the ones reported here have the potential to result in viruses in which the target of a PCR diagnostic test is deleted. Using a PCR assay that targets an essential gene (OPX3 or VAC1) or using multiple PCR assays that target different regions of the genome may be necessary to avoid identification of viruses with large deletions. PCR assays targeting non-essential genes in the terminal regions may be most at risk.

These unique genome rearrangements indicate that there is a continued need for genomic surveillance of MPXV to study its evolution during this outbreak and ensure current diagnostic approaches and medical treatments remain effective.

## Acknowledgements

The authors wish to thank Abby Berns and the other members of the Rhode Island Department of Health Center for Acute Infectious Diseases and Epidemiology team that are partners in the RI response to monkeypox and expertly organized patient testing and collected all demographic information; Nicholas Chen, Chantal Vogels, and the other members of the Grubaugh Lab in the Department of Epidemiology of Microbial Diseases, Yale School of Public Health, for the design of the MPXV tiled PCR-based amplicon protocol (<u>Monkeypox virus multiplexed PCR amplicon sequencing (PrimalSeq) V.2. (protocols.io)</u>; technical contributions that were made by the staff of the Maryland Department of Health Laboratory; and efforts made by the 2022 Multi-National Monkeypox Outbreak Response teams at the Centers for Disease Control and Prevention, Minnesota Department of Health, Florida Department of Health, DC Health, and Massachusetts Department of Public Health. T. Khan was supported in part by appointment to the Research Participation Program at the Centers for Disease Control and Prevention, administered by the Oak Ridge Institute for Science and Education through an interagency agreement between the U.S. Department of Energy and CDC.

## Materials and Methods

### Sequencing

#### ONT

Library preparation was performed on extracted DNA using Ligation Sequencing kit (Oxford Nanopore Technologies SQK-LSK-109) following the manufacturer’s protocol for genomic DNA. Libraries were sequenced (one sample per flow cell) on a MinION sequencer (RI050-7718) or GridION sequencer using a MIN109 R9.4.1 flow cell (Oxford Nanopore). Data are from a single swab from a single lesion. Basecalling was performed using guppy version 6.1.2 with high accuracy and qscore filtering (for MinION runs) or was performed on GridION using high accuracy basecalling.

#### Illumina

Extracted DNA (15 µL) was used as input for the Illumina DNA Prep method according to the recommended protocol with ½ reagent volumes used throughout. Libraries were visualized using the Agilent Fragment Analyzer instrument and the HS NGS Fragment Kit (Agilent Technologies Inc., Santa Clara, CA). Forty-eight libraries were pooled at approximately equal molarity and sequenced (200 pM final loading concentration) on an Illumina NovaSeq 6000 instrument using the 300 cycle SP sequencing components.

### Genome Assembly

Nanopore reads were trimmed to remove 55 bp from each end (seqtk 1.0, https://github.com/lh3/seqtk) and all reads below 50 bp were removed (trimmomatic 0.39, https://github.com/timflutre/trimmomatic) before mapping to MPXV Nigeria reference MT903344 with 6,000 bp removed from the left terminus using minimap2 2.16 (https://github.com/lh3/minimap2) to remove human and other non-MPXV reads. Illumina reads were trimmed using FaQCs 1.34 (https://github.com/LANL-Bioinformatics/FaQCs) using parameters -q 20 --5end 10 --3end 5 -n 5 --min_L 30 and then mapped to MPXV lineage B.1 reference sequence ON563414.3 using bwa mem (https://github.com/lh3/bwa). A hybrid assembly was generated from mapped Illumina and Oxford Nanopore reads using Unicyler 0.4.7 (https://github.com/rrwick/Unicycler). Assemblies were polished by mapping reads back to draft genomes containing one complete ITR and one incomplete ITR with ~6,000 bases removed using bwa mem or minimap2 and generating a consensus sequence using samtools 1.9 and ivar 1.0 (https://github.com/andersen-lab/ivar). Inverted Terminal Repeats (ITRs) were assembled manually by copying from one end to the other.

For quality control, separate assemblies were made with either nanopore or Illumina data since DNA extracted from separate lesions for the same patient were used for Nanopore and Illumina sequencing in three cases. Illumina-only assemblies were made using SPAdes/3.13.0 and CLC Genomics Workbench 22. Oxford Nanopore-only assemblies were made using flye 2.9 (https://github.com/fenderglass/Flye).

Annotations were transferred from MPXV Clade 3 Nigeria reference MT903345, then locus_tags were re-named with the strain ID. Initial alignments used to make trees were performed using MAFFT v.7.450 (*31*). To better visualize alignments, gaps were manually added to separate sequence from 5□ and 3□ ends of the genome. Then regions of the alignment were re-aligned using MAFFT.

**Table S1.**
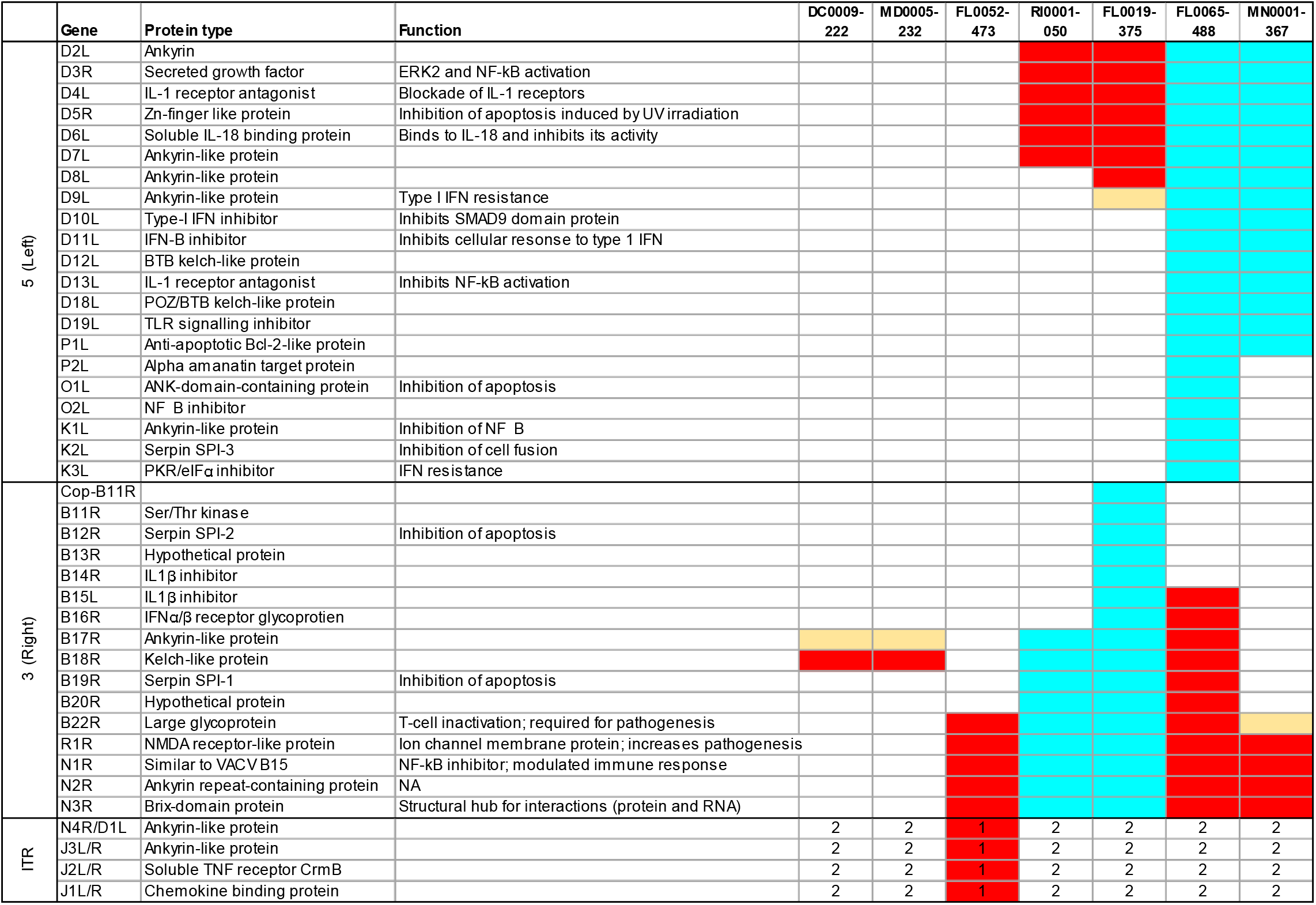
List of CDS duplicated or deleted in seven 2022 U.S. MPXV with large genomic rearrangements. Red: deleted; cyan: duplicated; yellow: predicted truncation. Copy number is given for CDS in ITR.

Text S1. Reads spanning rearrangement or deletion junction points.

**Figure S1.**
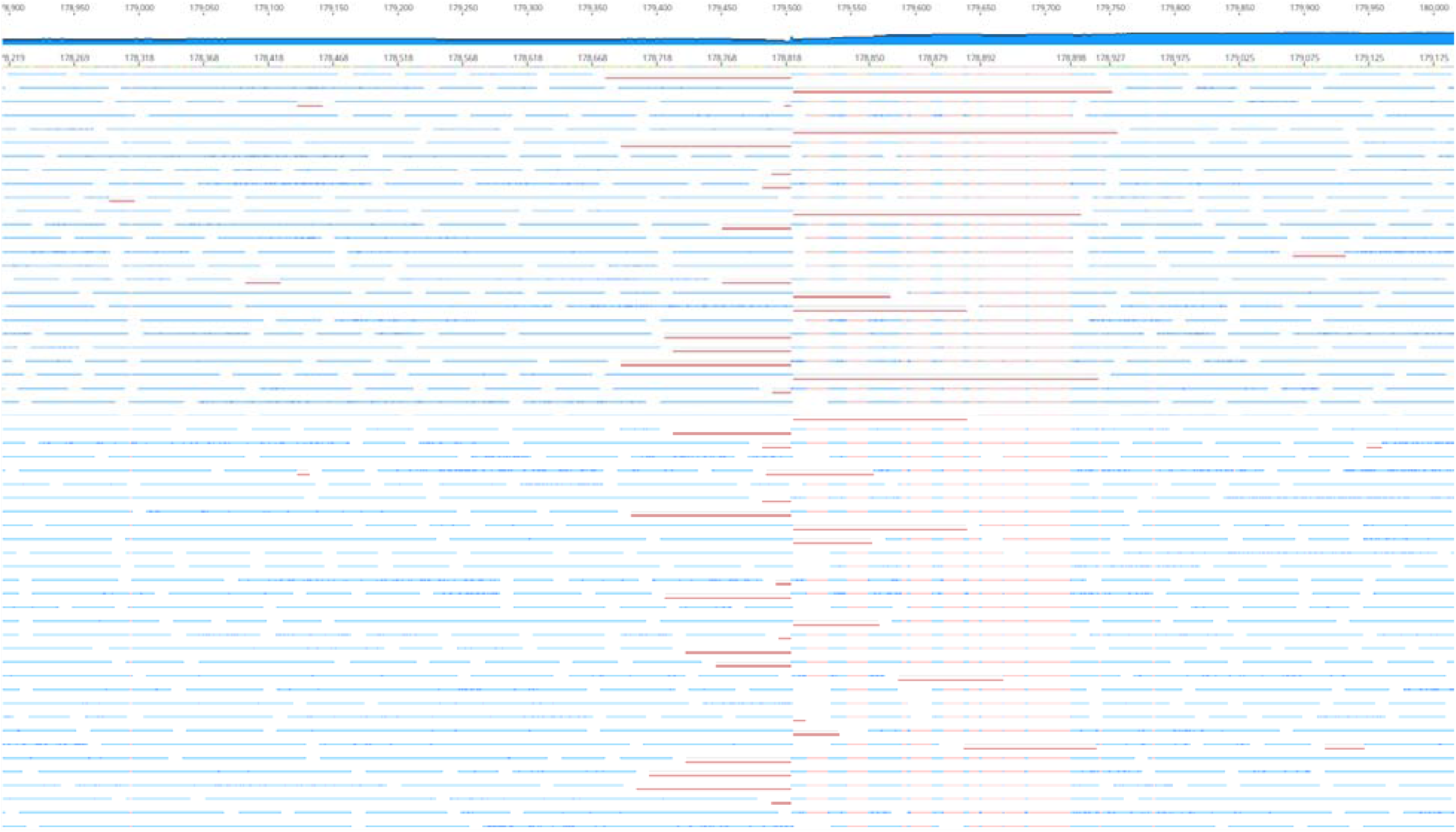
Reads mapped to test draft genome where no coverage region was converted to a simple deletion. Pink bars indicate masked portions of reads that contain sequence that does not match the reference. All reads that cross the hypothetical junction only map to one side or the other. No reads cross the junction, suggesting the simple deletion is not supported by the data.

**Figure S2.**
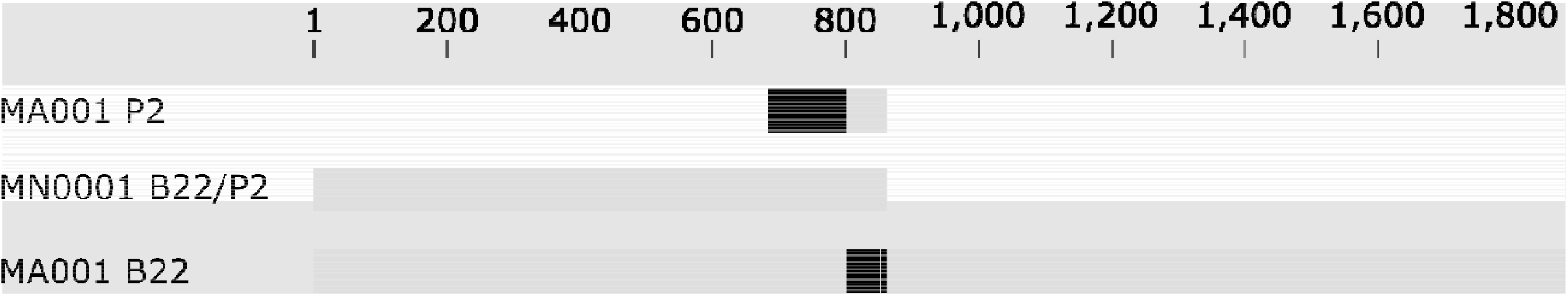
Predicted hybrid B22 protein in MN0001. Amino acid alignment of MN001 B22 protein compared to MA001. MN001 B22 protein is identical to MA001 B22 for the first 801 aa that precede the genomic rearrangement. The predicted MN0001 B22 protein then contains a 60 aa insertion, resulting in a predicted protein missing the C-terminal 1,019 aa of MA001 B22R. Gray bars indicate identity, black indicates differences.

**Figure S3.**
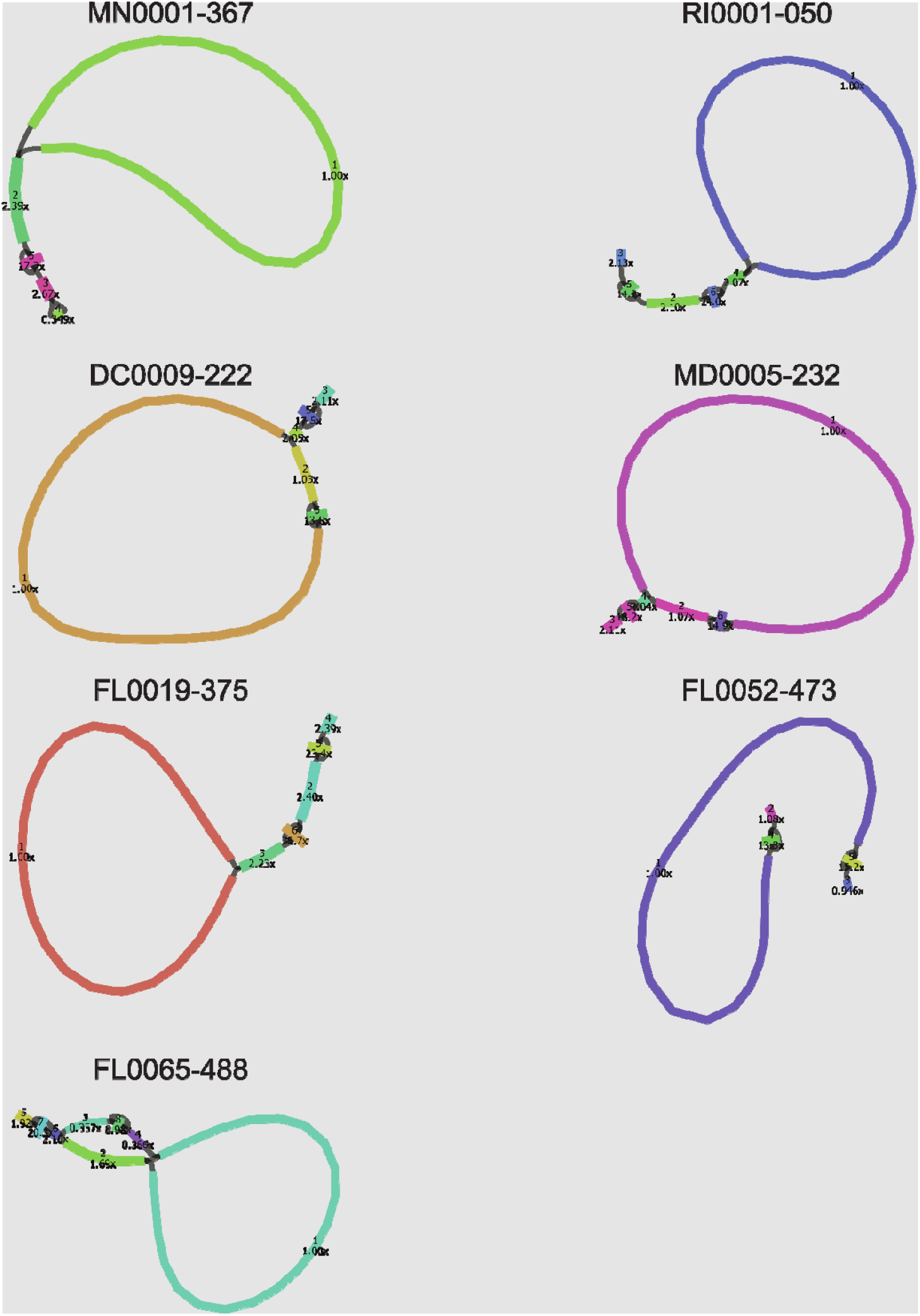
Model of contig assembly for seven MPXV genomes. Looped contigs are shown connecting to contigs with 2x or higher coverage at both ends. A tandem repeat region with in the ITR is shown with very high coverage compared to the main contig. Contig map was produced in Bandage from unicycler output data.

